# Size Matters: Small Cell Variants of *Coxiella burnetii* Initiate Replication Early in Primary Macrophages

**DOI:** 10.64898/2026.08.17.744959

**Authors:** Leslie A. Sims, Parker A. GrandPre, Shawna C. O. Reed, Elizabeth Di Russo Case

**Author notes:** These authors contributed equally to the manuscript (Leslie A. Sims, Parker A. GrandPre).

## Abstract

*Coxiella burnetii* alternates morphologies to survive in two niches: the external environment and a degradative intracellular compartment. The small cell variant (SCV) is adapted for environmental persistence and transmission of Q fever to ruminants and humans. The large cell variant (LCV) is intracellular, and despite not being a major source of transmission, is infectious *in vitro*. When modeling infection, researchers typically apply a mixed population of these cell types as inocula. As this practice does not mimic natural infection, it may confound our understanding of early Q fever infection events. We separated SCV and LCV by density gradient centrifugation and compared their replication in primary murine macrophages and a fibroblast cell line. SCV inocula replicated more efficiently than LCVs in both host cell types. LCV replication was delayed for four days in macrophages compared with SCV inocula, which had completed logarithmic growth by that time point. We found no difference in pathogenic vacuole size, but there was a modest difference in their respective bacterial burdens. Interestingly, IL-6 and CXCL2 secretion was significantly elevated in LCV-infected macrophages as compared to SCV at 24 hours, suggesting a difference in the host response to each. This is the first study to demonstrate that *C. burnetii* developmental status influences the progression of infection.

## Introduction

*Coxiella burnetii* is a Gram-negative, obligate intracellular bacterium and the highly infectious agent of the zoonotic disease Q fever [1,2]. It is environmentally resilient and can survive in the environment for several years [3–6] due to its documented resistance to UV radiation, desiccation, extreme temperatures, osmotic pressure, and chemical disinfectants [1,7]. It easily aerosolizes and transmits to the subsequent host through inhalation [8]. *Coxiella* can infect and replicate within various host cell types both *in vivo* and *in vitro*. However, alveolar macrophages remain the primary target cell in humans, and placental trophoblasts in ruminants [11].

*C. burnetii’s* biphasic lifestyle enables its environmental stability and transmissibility. The bacterium alternates between two morphologies: a small cell variant (SCV) and a large cell variant (LCV), which differentiate in response to environmental conditions [1,12,13]. The SCV is the metabolically inactive extracellular form optimized for transmission and environmental survival. Its compact structure has a thick, rigid cell wall enriched in lipopolysaccharide (LPS), which enhances its environmental resistance and enables immune evasion by masking immunogenic surface structures [14,15]. These properties make SCVs highly infectious and capable of infecting a new host when conditions favor replication, particularly at an acidic pH [1,16,17]. In contrast, LCVs are the metabolically active, replicating intracellular form that quickly lose viability outside a host [12]. A major hallmark of the transition from SCV to LCV is a shift in peptidoglycan structure from shorter, rigid 3-3 cross-links to 4-3 cross-links [18], resulting in a thinner, more flexible cell wall. Moreover, a shift in LPS expression exposes more surface proteins, making LCVs more antigenic [18–21].

The mechanisms that regulate differentiation between the cell variants, beyond the requirement for an acidic environment, remain mysterious [16]. However, during natural infection, the process is thought to begin when a new host cell engulfs a SCV. *Coxiella* is taken up by an immune cell and trafficked to a low-pH phagolysosome, where it transitions into a LCV [1]. The bacteria deliver effector proteins via the Type IVb Secretion System (T4SS) to transform their bounding vacuole into a pathogen-tailored compartment known as the *Coxiella*-containing vacuole (CCV), with an approximate pH of 5 [12,22,23]. *Coxiella* replicates within this vacuole, and once the replication cycle is complete, LCVs differentiate back into SCVs in response to as-yet-unidentified signals [10,13,19,20,24].

Research indicates that both SCVs and LCVs are similarly infectious *in vitro* across various cell types [12,25,26], regardless of tissue origin or species [27,28]. Natural infections similarly show that a variety of cell types can support bacterial growth, including splenocytes, hepatocytes, Kupffer cells in the liver, and adipocytes [29–31]. However, alveolar macrophages are considered the primary natural host targets in humans, and placental trophoblasts in infected ruminants [32,33]. Wiebe’s landmark study in 1972 used density gradient centrifugation to physically separate SCVs and LCVs and compared their viability in embryonated hen’s eggs [25]. The isolated LCVs were shown to be approximately 15-fold more lethal and 5-fold more infectious than the SCVs. While this study established that both cell variants are infectious, the techniques used were of limited sensitivity and lacked relevance to natural infection. For example, infectious doses were determined using optical density, a technique that is inaccurate for enumerating *C. burnetii* inocula [34].

Recent technological advances in molecular and cellular approaches for studying *C. burnetii*-host interactions now allow researchers to revisit this question with greater precision. In 2009, the development of an axenic culture medium enabled the cultivation of *Coxiella* without the use of animals or cells. Additionally, qPCR and physiologically relevant cellular models [35–37] have greatly enhanced our understanding of *C. burnetii* physiology during infection. These tools, combined with density gradient centrifugation, enable a more direct comparison of cell variant-dependent replication kinetics and host immune responses, thereby clarifying their distinct roles in infection.

In this manuscript, we compared the replication phenotypes of density gradient-purified SCVs and LCVs in primary murine macrophages with those in a stable fibroblast cell line that is commonly used to model *C. burnetii* infection [35,38,39]. We hypothesized that LCVs, due to their fragility and highly immunogenic nature, are ill-adapted to establish infections in macrophages, as compared to SCVs. Indeed, SCVs initiated replication earlier in primary macrophages than LCVs, which exhibited a prolonged lag phase. We also observed that the generation time of each cell variant was significantly shorter in BMDMs than in L929 cells. Surprisingly, while the SCV and LCV inocula exhibited similar lag phases in L929 murine fibroblasts, the bacteria from the SCV infections replicated faster in these cells compared to LCV infections. These stark differences in replication kinetics highlight the contribution of both the host cellular environment and *C. burnetii*’s developmental status to infection progression. We also investigated host inflammatory responses to infection with SCVs and LCVs. We observed modest differences in the host response to both infecting cell variants at 24 hours post-infection. This suggests that macrophages may sense and respond to the two cell variants differentially; however, we cannot assume these responses are the sole factor influencing *C. burnetii* replication kinetics.

## Materials and Methods

### Bacteria

This study used the *Coxiella burnetii* Nine Mile, Phase II, clone 4, plaque-purified strain. This strain is excluded from the National Select Agent Registry under HHS regulations (42 C.F.R. Part 73) and is approved for use under Biosafety Level 2 containment. In compliance with the University of Wyoming Institutional Biosafety Committee, the strain’s lack of *cbu0533* gene reversion was verified prior to the initiation of this work. All cultures were grown in acidified citrate cysteine medium-2 (ACCM-2; Sunrise Science Products) [29] from a single frozen stock. Approximately 10 μL of the stock was used to inoculate 1 L of medium. Cultures were incubated undisturbed in horizontally oriented vented tissue culture flasks at 37 °C with 5% CO_2_ and 2.5% O_2_ for 7 days.

### Cell Culture

Murine L929 fibroblasts (ATCC) were cultured in complete DMEM, consisting of Dulbecco’s Modified Eagle Medium (DMEM) supplemented with 10% (v/v) heat-inactivated fetal bovine serum (FBS). Primary bone marrow-derived macrophages (BMDMs) were differentiated from the bone marrow of C57BL/6 mice (The Jackson Laboratory) as previously described [40]. The BMDMs were also maintained in complete DMEM supplemented with 20% (v/v) L929 cell-conditioned medium (LCCM) until the day of infection. Thereafter, the LCCM was maintained at 15%.

### Density Gradient Centrifugation

To increase bacterial yield, *C. burnetii* was first cultured in 5 mL of ACCM-2 for 7 days, then passaged into 1 L of fresh ACCM-2 for an additional 7 days. After incubation, the bacteria were harvested by centrifugation at 16,000 x *g* for 16 minutes at 4 °C. The pellet was washed in 20 mL of cold, sterile 0.25 M Sucrose Phosphate (SP) buffer [41] and resuspended in 48 mL of SP buffer. The bacterial suspension was distributed into 6 tubes layered on top of 22 mL of SP buffer and 8mL of 30% Gastrografin. The tubes were centrifuged at 58,400 x *g* at 4 °C for 30 minutes. Following the centrifugation, the supernatant was removed, and the bacterial pellet was resuspended in 50 mL of fresh SP buffer and incubated overnight at 4 °C.

Density gradients were prepared in ultra-clear, open-top, thin-walled centrifuge tubes (Beckman Coulter). A discontinuous gradient was prepared by mixing cold SP buffer with increasing concentrations of Gastrografin (Spectrum Medical Imaging Co.) to final concentrations of 54%, 44%, and 40% (v/v). Gradients were layered in the centrifugation tubes in the following order, from bottom to top: 8 mL of 54% solution, 12 mL of 44% solution, and 8 mL of 40% solution. 10 mL of the bacterial sample in SP buffer was layered on top of each gradient. Centrifugation was conducted at 58,420 x *g* for 1 hour at 4 °C [42]. SCVs formed a band above the 40-44% interface, and LCVs below the 44-54% interface. Each band was collected by piercing the tubes with a sterile 18-gauge needle with the needle bevel facing up. The bands were collected with an attached 10mL syringe and then transferred to separate sterile Oakridge centrifuge tubes.

Each tube containing the extracted bacteria was adjusted to a final volume of 25 mL with SP buffer. The bacteria were pelleted and then washed once with 10 mL of SP buffer, as described above. The final pellets were resuspended in SP buffer to approximately 1 x 10^7^ cells/μL and stored at −70 °C until used in downstream assays. The total concentration of *C. burnetii* in each stock was determined using a TaqMan qPCR assay targeting the *com1* gene [43]. Genomic DNA from each stock was extracted in triplicate, and each sample was amplified in 3 technical replicates.

### Viability Assay

Bacterial viability was assessed by spot plating on plates containing 1X ACCM-2 and 0.5% Ultra-Pure Agarose (Thermo Scientific). *C. burnetii* stocks were serially diluted from 10^-3^ to 10^-8^ in 2X ACCM-2, and an equal volume of 0.5% agarose was mixed with the diluted bacteria for a final concentration of 0.25% for spotting. 50 μL of each dilution was spotted in triplicate onto the prepared ACCM-2 plates and incubated for 10 days. Colonies were then counted under a light microscope.

### *In vitro* Infection Assays

All *in vitro* infections used both murine L929 fibroblasts and murine bone marrow-derived macrophages (BMDMs), maintained as described above. For BMDMs, a different mouse was used to derive the macrophages for each infection. Infections were conducted as previously described [37], with the modifications outlined below. A single inoculum was prepared for either SCV or LCV bacterial populations and applied to L929 cells and BMDMs in parallel infections at a multiplicity of infection (MOI) of 25 bacteria per host cell. The culture medium was replaced daily, and cells were monitored by microscopy for health.

For each infection condition, triplicate wells were processed at the following time points: days 0, 1, 4, 7, and 10 post-infection. Day 0 samples were collected immediately after the inoculum was washed off the cells at 1-hour post-infection. At each time point, the medium was replaced with 200 μL of lysis buffer [43]. Lysates from each well were transferred into separate microcentrifuge tubes for genomic DNA extraction and bacterial enumeration, as described above. Generation times were calculated from changes in genome equivalents during the log phase of bacterial growth for each condition. In BMDMs, this was defined as days 1-4 for SCV infections, and days 4-7 for LCV infections. In L929 cells, days 1-7 were used for both. The data presented are the mean of 3 independent infections, and statistical significance was determined using 2-way RM ANOVA followed by Šídák’s multiple comparisons test.

### Antibody Generation

A rabbit polyclonal anti-ScvA antibody was custom generated by Genscript, Inc. against a synthetic polypeptide corresponding to the full 30-amino acid sequence of the ScvA protein with CM at the N-terminus [CMERQNVQQQRGKDQRPQRPGASNPRRPNQR]. The antibody was affinity purified against the synthetic peptide and validated for specificity using ELISA in comparison with non-specific IgG.

### Western Blot Analysis

Western blotting was used to compare the relative expression of the SCV marker protein ScvA across bacterial populations isolated by gradient centrifugation. Whole-cell lysates containing 1 x 10^7^ bacteria/μL, as determined by qPCR, were prepared according to the manufacturer’s instructions using the 4x Protein Sample Loading Buffer (LI-COR), and then boiled for 10 minutes. The equivalent of 1 x 10^8^ bacteria were loaded per lane and separated using 4-20% gradient SDS-PAGE (Bio-Rad) and NuPAGE MES SDS Running Buffer. The primary antibodies used were rabbit anti-Com1 (gift of Dr James Samuel, Texas A&M University Health Science Center) and rabbit anti-ScvA. The membranes were imaged using the IRDye® 680RD Donkey anti-Rabbit Antibody (LI-COR) on the LI-COR Odyssey XF, with the 700- and 800-nm channels exposed for 2 minutes each.

### Protein Secretion Array

BMDM culture supernatants were collected immediately prior to infection and again at 24 hours post-infection, when the cells were fed. At each time point, supernatants from the same 3 replicate wells were pooled into a 2 mL microcentrifuge tube and centrifuged at maximum speed for 10 minutes to pellet any cellular debris. The supernatant was carefully removed and transferred into a new microcentrifuge tube for storage at −70 °C. Cytokine arrays were performed according to the manufacturer’s instructions for the Proteome Profiler Mouse XL Cytokine Array (R&D Systems), including their supplemental protocol for imaging on the LI-COR Odyssey system.

Cytokine arrays were performed on 3 independent infections using BMDMs isolated from 3 individual mice. Data were processed by subtracting background from the signal, averaging the signal from each of the two replicate spots on the array, and calculating the fold change in signal for each infected sample relative to its matched uninfected control supernatant. The data presented are the mean fold changes from 3 independent experiments; error bars represent standard deviations, and statistical significance was determined using 2-way RM ANOVA followed by Uncorrected Fisher’s Least Significant Differences test. PCA analysis was conducted on background-subtracted signals for all conditions using Cytoprofile (https://cytoprofile.cytokineprofile.org/) [44].

### Fluorescence Microscopy

Coverslips were prepared as described previously [37] and collected at the indicated time points. Coverslips were then stained, with *Coxiella* detected by primary guinea pig NMII antiserum (gift of Dr James Samuel), rat α-LAMP1 antibody (DSHB, University of Iowa), and 100 ng/ml of DAPI (4’,6-diamidino-2-phenylindole). Stained coverslips were mounted to a microscope slide using MOWIOL.

Images were captured using a Leica Stellaris 8 FALCON FLIM-STED microscope with a 63x oil immersion lens. Images were processed using the proprietary LAS X software. CCV size was quantified using the Fiji distribution of ImageJ. Mean CCV areas were compared by 2-way RM ANOVA followed by Bonferroni’s multiple comparisons test, and no statistical significance was found.

To determine bacterial burdens in individual CCVs, bacteria from at least 100 CCVs were counted per condition across the 3 independent experiments using an Olympus BX51 fluorescence microscope with a 63X oil-immersion lens. Data analysis using the Kruscal-Wallis test followed by Dunn’s Multiple Comparisons test revealed no statistical significance.

## Results and Discussion

### Separation of SCV and LCV by Discontinuous Gradient Sedimentation

Our aim for this study was to determine whether SCVs and LCVs differ in their ability to establish an infection *in vitro*. This required the physical separation of the two cell types (Fig 1A). We opted to use axenic (cell-free) culture to raise our bacterial inocula for several important reasons. First, this approach allowed us to assess the sufficiency of each cell type for infection without indirect effects from contaminating host material. Second, as multiple studies have shown, a 7-day ACCM-2 culture contains mixed populations of SCVs and LCVs [18,24]. Therefore, isolating both variants at a single time point eliminated confounding effects from differing ages or culture conditions. Finally, culturing bacteria in broth eliminated labor-intensive and time-consuming isolation steps [26], minimizing bacterial stress and maximizing viability. After one hour of centrifugation through a discontinuous Gastrografin density gradient, the *C. burnetii* culture separated into three distinct bands, each representing a bacterial population with a different buoyant density (Fig 1B). The bands were extracted sequentially from top to bottom using a syringe fitted with an 18G needle. The first band was at the bottom of the 40% Gastrografin layer, the second at the top of the 44% layer, and the third at the top of the 54% layer of the step gradient.

**FIG 1.**
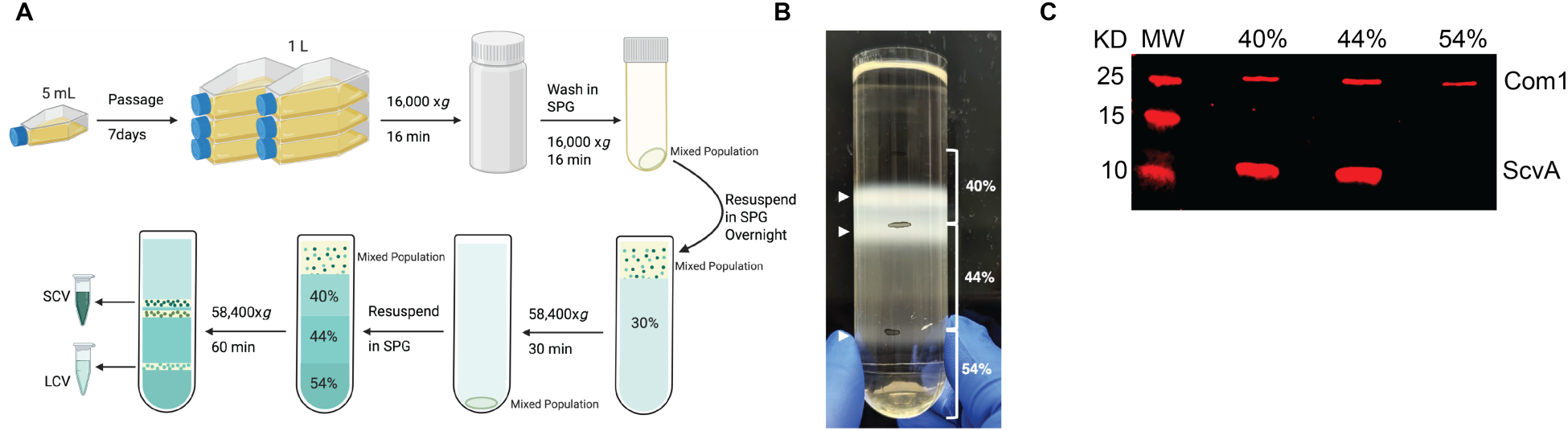
Separation of SCVs and LCVs by density gradient. (A) Workflow illustrating the separation of SCVs and LCVs via discontinuous gradient sedimentation. Created in BioRender. Case, E. (2026) https://BioRender.com/5xmhmlq (B) Fractionation of NMII cell types results in three distinct layers at the 40%, 44%, and 54% Gastrografin layer boundaries. (C) Western blot confirming ScvA expression phenotypes for each of the separated bacterial populations. Mouse anti-ScvA was used as a marker for SCVs. The constitutively expressed Com1 protein was used as a loading control.

We note that we could find no other description in the literature of a density gradient separation for *C. burnetii* that resulted in three bacterial populations [25,45–48] In our hands, the band was only evident after overnight equilibration in SP buffer at 4°C. Other methodological differences between our study and previous studies may also contribute to this observation. To our knowledge, this is the first study to perform density gradient separation using axenically cultured bacteria. Axenic cultivation enabled the recovery of substantially larger quantities of bacteria without the need for host-cell propagation and subsequent purification, potentially influencing the composition of bacterial populations subjected to density gradient centrifugation.

We next used western blotting to determine which of the isolated populations expressed the SCV marker protein ScvA [12,47]. We loaded 1 x 10^8^ bacteria per lane and probed for the constitutively expressed Com1 protein as a loading control. Both bacterial populations isolated from the 40% and 44% Gastrografin layers expressed ScvA, whereas the third population at the 54% layer showed very little detectable ScvA protein compared with the Com1 control (Fig 1C). Based on this, we defined the bacterial population isolated from the 40% Gastrografin layer as SCVs and that from the 54% layer as LCVs, according to their ScvA expression phenotypes [25,45–48] and respective densities. The isolation of SCVs from the uppermost layer of the Gastrografin gradient is consistent with previous data in which the LCV forms increase in apparent density due to diffusion of the gradient medium across their membranes [1,25,47]. We elected not to perform downstream experiments with the bacteria isolated from the 44% Gastrografin layer, as we could not be assured of the population’s homogeneity. It is possible that the population included transitional forms of intermediate density, though this requires further investigation.

Prior to infection, it was important to determine the relative viability of the recovered bacterial populations. We first determined the viable counts of the bacteria isolated from each band by plating them on solid ACCM-2. For the 40%, 44% and 54% bands, we recovered a total of 2.83 × 10^10^, 4.90 × 10^10^, and 2.82 × 10^9^ bacteria, respectively. Across multiple independent gradients, bacterial abundance in the upper bands was consistently higher than that of the 54% Gastrografin population. By enumerating total bacteria with qPCR, we found that SCVs exhibited 47.94% viability, whereas LCVs showed 17.52%. We attributed the approximate 2.7-fold difference in viability to the osmotic stress LCVs experience during passage through the density gradient.

### SCV and LCV Exhibit Distinct Intracellular Replication Phenotypes

After successfully separating the SCV and LCV populations from a single culture, we sought to test their ability to establish *in vitro* infection in murine fibroblasts and bone marrow-derived macrophages. L929 fibroblasts were selected for their established permissiveness to *C. burnetii* replication [35,38,39,49–51]. BMDMs were chosen as a comparatively restrictive host cell type that has only recently been established for use with *C. burnetii*, Nine Mile Phase II [36,37]. We reasoned that comparing these two cell types, which differ in their ability to support *C. burnetii* replication, would maximize any measurable differences in infectivity between the developmental variants.

We infected both cell types in parallel infections using the same inocula. At 0, 1, 4, and 7 days post-infection, whole cell lysates were collected for bacterial enumeration by qPCR. We also collected genomic DNA from infected BMDMs after 10 days, but we found it difficult to maintain the L929 cells in culture for that length of time. As shown in Fig 2, the replication kinetics of the SCV and LCV inocula differed in both host cell types. In the L929 cells, both SCV and LCV inocula exhibited lag phases of comparable length (Fig 2B). However, generation time was longer (Fig 2C) in cells infected with LCVs, averaging 34.8 hours, compared to 21.8 hours for infections initiated with SCVs. In contrast, the SCV inocula replicated rapidly in BMDMs from days 1 to 4 post-infection before entering stationary phase. When infection was initiated with LCVs, *C. burnetii* exhibited a protracted lag phase in these same host cells, entering the logarithmic growth phase on day 4 (Fig 2A). We found this surprising because LCVs do not require a developmental transition prior to replication, as SCVs do. Intriguingly, the generation time for both inocula was significantly shorter in BMDMs than in fibroblasts (Fig 2C), suggesting that the CCVs established within BMDMs supported more efficient replication than those of L929 cells. The mean generation time for LCV inocula was 20.3 hours, and for SCV, 12.6 hours. Comparing the generation time differences across conditions revealed that SCV inocula sustained a replication rate that was 1.6-fold faster than LCV in both host cell types. This is the first demonstration that *Coxiella burnetii* exhibits a developmental difference in replication kinetics.

**FIG 2.**
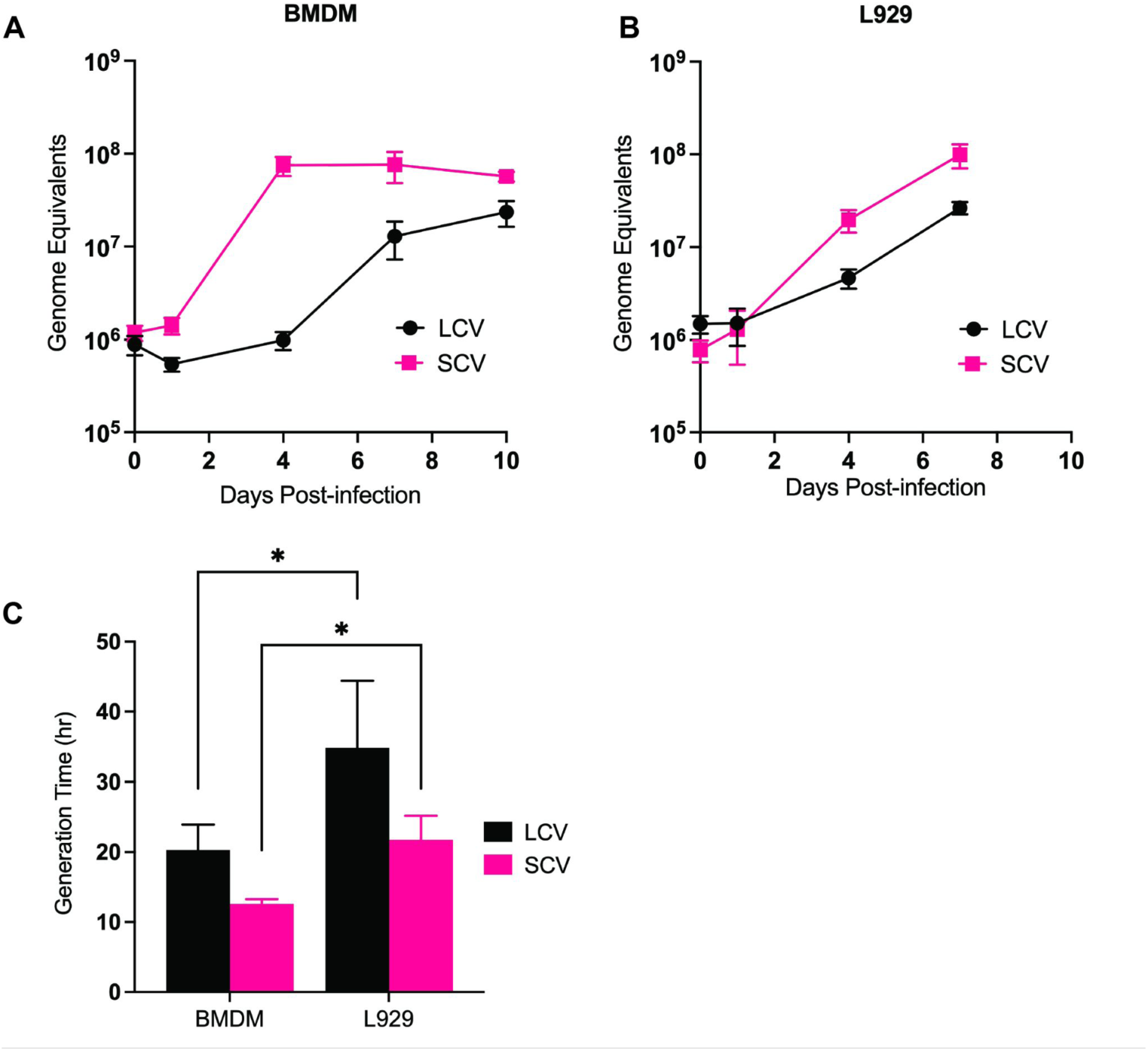
Intracellular replication of SCVs and LCVs in BMDMs and L929 cells. (A and B) Growth curves of SCVs and LCVs grown in BMDMs (A) over 10 days or L929s (B) over seven days. Each data point is the mean from 3 independent experiments. Error bars represent standard deviations. (C) Generation time of SCVs and LCVs grown in either BMDMs or L929s. Data were calculated from the logarithmic growth periods: days 1 to 4 for SCVs, and days 4 to 7 for LCVs in BMDMs, days 1-7 in L929 cells. Data are the mean of 3 independent experiments, with error bars indicating standard deviations. Asterisks represent *P-*values determined by Two-Way RM ANOVA, using Šidák’s multiple comparisons (*, *P=*0.0391).

To investigate differences in CCV size in BMDM infected with either SCVs or LCVs, we imaged coverslips collected at 4 days post-infection. We focused on the CCV phenotypes in BMDM as this was the condition with the greatest observed difference in bacterial replication. In general, CCV size did not differ significantly between cells infected with SCVs or LCVs (Fig. 3C). Indeed, the range of CCV areas we measured varied widely in both conditions, both within and across experimental replicates. We then assessed the bacterial burden in BMDMs infected with LCVs compared with SCV inocula. Counting the bacteria within CCVs of infected BMDMs, we found that LCV-infected monolayers had proportionally fewer CCVs containing greater than 5 bacteria compared to SCV-infected cells (Fig. 3B). While the differences in the bacterial burden of individual CCVs were not statistically significant, we believe that the proportion of CCVs with greater than 100 bacteria in the SCV-infected BMDM likely contributed to the difference in total bacterial numbers measured at 4 days post-infection (Fig 2).

**FIG 3.**
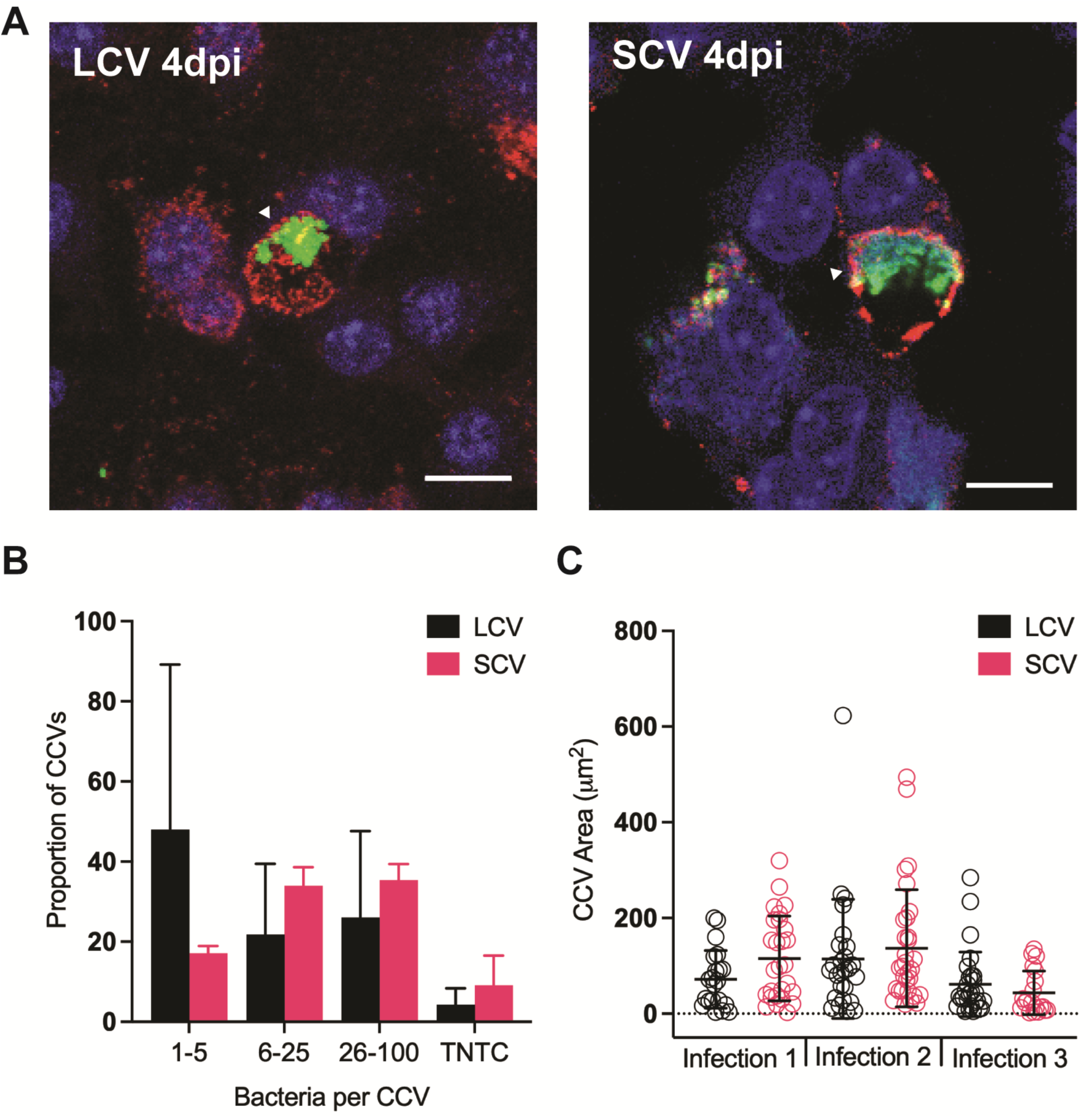
CCV size and bacterial burden in BMDMs infected with LCVs or SCVs. (A) Representative images of CCVs (arrowheads) from BMDMs infected with LCVs (left) or SCVs (right) at 4 days post-infection. Bacteria are stained green, LAMP1 is red, and DNA is blue. Scale bar = 10 μm. (B) Bacteria were enumerated to determine burden in each of 100 CCVs. Data are the mean of 3 independent experiments, with error bars indicating standard deviations. (C) Cross-sectional areas of CCVs were measured in μm^2^ using the Fiji distribution of ImageJ. Each point represents the area of single CCV. Error bars represent standard deviations.

### Both *C. burnetii* Cell Types Induced Modest Protein Secretion in Macrophages

It is important to note that whether we initiate an *in vitro* infection with SCV or LCV inocula, all intracellular replication is carried out by bacteria in the large cell variant form. These data suggest a fundamental difference in replication progression depending on which cell variant initiates infection. At the outset of this study, we hypothesized that because of the SCVs’ extraordinary environmental stability and low antigenicity [14,15], they are better adapted to invade and survive within a macrophage than the LCV bacterial form. Conversely, we posited that, as LCVs have a comparatively fragile structure and are more antigenic [45] they may have reduced stealth, stability, and therefore, infectious potential. To address this possibility, we next sought to determine if the differences in bacterial replication we observed in BMDMs corresponded to a differential innate immune response in the host. To do this, we applied cell culture supernatants from 24 hours post-infection to a protein secretion array (Fig 4). The array allowed for qualitative comparison of proteins secreted by infected and uninfected BMDMs. We elected to assess protein secretion early, before the SCV-to-LCV transition could occur, reasoning that this would best capture the immediate host response to either cell type. We observed that upon infection, multiple host cytokines and chemokines were upregulated by both SCV and LCV populations compared with uninfected cells. Principal Component Analysis revealed that SCV-infected cells showed greater variance in the level of cytokine secretion than those infected with LCVs (Fig 4A).

**FIG 4.**
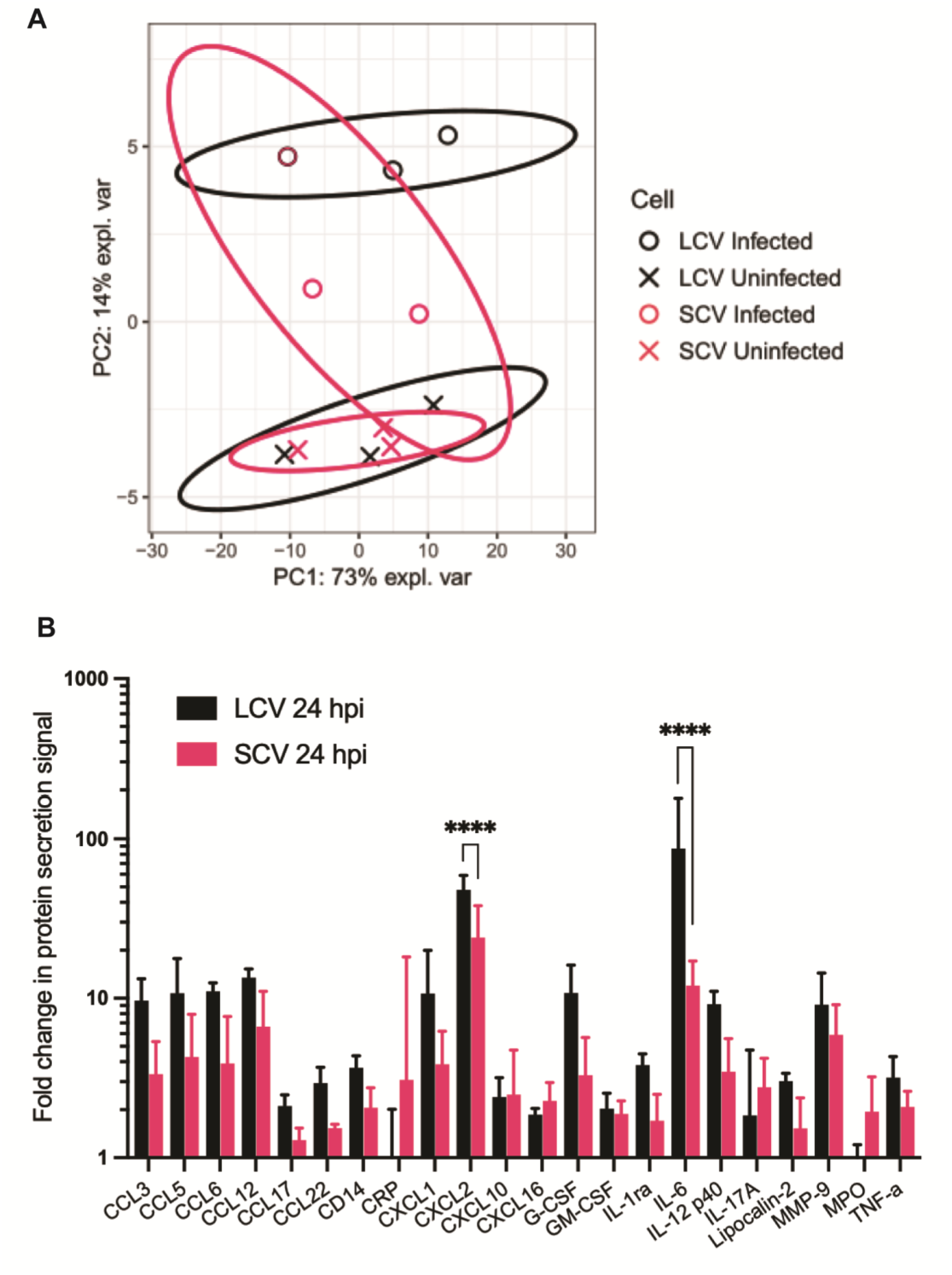
Protein secretion by BMDMs infected with LCVs or SCVs. (A) Principal Component Analysis of secretion profiles of uninfected BMDMs or BMDMs infected with either SCVs or LCVs. Each point represents an individual experiment. Ellipses represent 95% confidence intervals of each condition. (B) Relative protein secretion in BMDMs infected with LCVs or SCVs. Shown are only proteins with >2-fold change in secretion over uninfected controls. Statistics were performed using Two-Way RM ANOVA, followed by Uncorrected Fishers Least Significant Differences test (****, *P*<0.0001).

Overall, infection with each cell variant induced a modest increase in cytokine secretion, with only 22 of 111 proteins tested showing a two-fold or greater increase compared with uninfected cells (Fig 4B). We observed no proteins that exhibited a 2-fold or greater decrease in secretion (Table S1). The identities of the upregulated proteins were largely consistent between variants and were predominantly chemokines involved in immune cell recruitment. Several key pro-inflammatory cytokines showed only modest induction. Secretion of TNF-α and IL-17A increased approximately two-fold for BMDMs infected with either SCVs or LCVs. Νο secretion of interleukin-1β (IL-1β) was detected above background levels (Table S1). There were two secreted proteins that distinguished infection with the two cell variants, IL-6 and CXCL2, both of which were secreted at higher levels in BMDMs infected with LCVs. IL-6 represented the most striking variant-specific difference, with 104.3-fold induction by LCVs compared with 12.5-fold in SCV-infected cells (Fig 4B). Secretion of CXCL2 was upregulated 48.6-fold in LCV-infected cells versus 25.9-fold in BMDMs infected with SCVs. While the number of differences we observed in protein secretion is limited, they are significant. This suggests that there may be differences in BMDM sensing and responses to the small and large cell variants of *C. burnetii*.

IL-6 secretion by *C. burnetii*-infected macrophages has been well documented, particularly in infection models that restrict *C. burnetii* replication [52]. IL-6 secretion in these cases has been attributed to detection of bacteria by TLR2 and TLR4. If the initial restriction of LCV-inocula we have observed in BMDM is TLR4-dependent, then this is consistent with the observation that the large cell variant expresses less LPS on its surface than its SCV counterpart [14,21]. The reduction in LPS expression has been shown to increase exposure of surface antigens and TLR2 ligands, while also reducing the TLR4-antagonistic effects of lipid A. The increased CXCL2 secretion in the context of LCV infection is less clear. A previous study documented that *C. burnetii* actively suppressed CXCL2 in a T4SS-dependent manner as part of the IL-17 pathway [53]. We did observe modest secretion of IL-17A and TNF-α in both infected cell populations; however, there was no significant difference in their secretion by cells infected with SCVs or LCVs. However, because the TLR4 pathway is also known to induce CXCL2 secretion, as with IL-6, this observation may reflect differences in LPS expression.

While these differences in cytokine secretion were interesting, we do not suspect that IL-6 and CXCL2 are the sole determinants of the observed developmentally-dependent replication kinetics. We note that even the permissive L929 cell line revealed a difference in *C. burnetii* replication rate between the two cell variants (Fig 2). We find it especially interesting that the SCVs, once transitioned to LCVs, maintain a consistently higher replication rate than the LCV inocula, independent of the host cell type infected. We believe this points to a developmental effect on the establishment of a replicative niche in the target host cell. In fact, a major functional distinction between SCVs and LCVs is their ability to secrete effectors by the T4SS. This secretion is essential for CCV development and replication. Perhaps effector secretion is sequentially ordered for proper CCV biogenesis after the SCV-to-LCV transition, and initiating infection with an LCV disrupts that temporal order. This could affect CCV development. There is data to suggest that T4SS effectors are temporally regulated. Gene expression data from Sandoz, et al. indicated several distinct developmental transcription profiles of known effectors. For example, *ankA* and *ankP* were transcribed at early times post-infection, while transcripts for *cirB* and *cig37* were detected late [18]. If developmental regulation of T4SS effectors influences CCV biogenesis, it is not an absolute requirement; LCV inocula ultimately are capable of replication. Future work will examine whether the characteristics of nascent CCVs differ when infections are initiated with either cell variant. It will be important to characterize these developmental effects on infection progression if the field is to understand how *C. burnetii* establishes a natural infection, which is typically initiated by the SCV.

## Supplementary Materials

<u>Table S1. Cytokine Array Data Summary</u>

## Author Contributions

Conceptualization, L.A.S., P.A.G., and E.D.C.; methodology, L.A.S., P.A.G., and E.D.C; formal analysis, L.A.S., and P.A.G..; investigation, L.A.S., P.A.G.; resources, E.D.C. and S.C.O.R.; writing—original draft preparation, L.A.S., P.A.G., E.D.C. and S.C.O.R.; writing—review and editing, E.D.C. and S.C.O.R.; visualization, P.A.G., and E.D.C; supervision, E.D.C. and S.C.O.R.; funding acquisition, E.D.C. and S.C.O.R. All authors have read and agreed to the published version of the manuscript.

## Institutional Review Board Statement

All experiments were conducted in compliance with the University of Wyoming Institutional Biosafety Committee (protocol number 2026-04-00, approved 11/17/2023). Likewise, the animal use protocol was approved by the University of Wyoming Institutional Animal Care and Use Committee (protocol number 2022-0054 and approved 03/22/2024).

## Informed Consent Statement

Not applicable.

## Data Availability Statement

The original contributions presented in this study are included in the article/supplementary material. Further inquiries can be directed to the corresponding author(s).

## Acknowledgements

The authors want to express their gratitude to Drs. Erin van Schaik and James E. Samuel for sharing reagents and helpful discussions. We want to thank Ben Sims and Cassi GrandPre for helpful discussions and for critically reading the manuscript. The authors acknowledge Brandoch Cook and Qian Yang at the Center for Advanced Scientific Instrumentation (CASI) at the University of Wyoming for training on and use of the "Cloud Peak" Leica Stellaris 8 FALCON FLIM-STED.

## Abbreviations

The following abbreviations are used in this manuscript

ACCM: Acidified citrate cysteine medium
BMDM: Bone marrow-derived macrophages
CCV: *Coxiella*-containing vacuole
DMEM: Dulbecco’s modified eagle medium
FBS: Fetal bovine serum
GE: Genome equivalent
LCCM: L929 cell-conditioned medium
LCV: Large cell variant
LPS: Lipopolysaccharides
MOI: Multiplicity of infection
SCV: Small cell variant
SP: Sucrose Phosphate
T4SS: Type IVb Secretion System

