## Supplemental Table 1 for "Size Matters: Small Cell Variants of *Coxiella burnetii* Initiate Replication Early in Primary Macrophages"

| | LCV uninfected Supernatants | | LCV Infected Supernatants, 24 hr | | SCV Uninfected Supernatants | | SCV Infected Supernatants, 24 hr | |
|---|---|---|---|---|---|---|---|---|
| Secreted Protein | Mean Signal | Standard Deviation | Mean Signal | Standard Deviation | Mean Signal | Standard Deviation | Mean Signal | Standard Deviation |
| adiponectin | 0.387 | 0.101 | 0.402 | 0.169 | 0.349 | 0.106 | 0.497 | 0.29 |
| amphiregulin | 0.561 | 0.171 | 0.414 | 0.313 | 0.584 | 0.304 | 0.502 | 0.315 |
| angiopoetin-1 | 0.628 | 0.17 | 0.402 | 0.197 | 0.487 | 0.083 | 0.443 | 0.225 |
| angiopoetin-2 | 0.512 | 0.188 | 0.314 | 0.197 | 0.362 | 0.161 | 0.222 | 0.193 |
| angiopoetin-like 3 | 0.563 | 0.221 | 0.343 | 0.183 | 0.39 | 0.134 | 0.282 | 0.248 |
| BAFF | 0.262 | 0.13 | 0.168 | 0.101 | 0.195 | 0.033 | 0.133 | 0.126 |
| C1qR1 | 0.441 | 0.125 | 0.375 | 0.181 | 0.327 | 0.12 | 0.254 | 0.169 |
| CCL2 | 3.177 | 0.455 | 3.85 | 2.115 | 3.34 | 1.632 | 2.623 | 1.879 |
| CCL3 | 0.405 | 0.093 | 3.244 | 1.504 | 0.38 | 0.121 | 1.036 | 0.425 |
| CCL5 | 0.257 | 0.052 | 2.847 | 1.582 | 0.402 | 0.141 | 1.163 | 0.212 |
| CCL6 | 1.247 | 0.08 | 11.148 | 5.938 | 0.961 | 0.276 | 3.101 | 1.356 |
| CCL11 | 0.531 | 0.169 | 0.366 | 0.238 | 0.462 | 0.074 | 0.32 | 0.141 |
| CCL12 | 1.053 | 0.073 | 11.747 | 2.124 | 0.874 | 0.187 | 5.914 | 3.133 |
| CCL17 | 0.722 | 0.205 | 1.047 | 0.7 | 0.442 | 0.192 | 0.49 | 0.271 |
| CCL19 | 0.573 | 0.182 | 0.393 | 0.243 | 0.378 | 0.175 | 0.338 | 0.311 |
| CCL20 | 0.189 | 0.05 | 0.134 | 0.072 | 0.118 | 0.051 | 0.087 | 0.064 |
| CCL21 | 0.35 | 0.081 | 0.26 | 0.145 | 0.232 | 0.107 | 0.156 | 0.1 |
| CCL22 | 0.567 | 0.139 | 1.218 | 0.843 | 0.478 | 0.223 | 0.624 | 0.292 |
| CD14 | 0.428 | 0.071 | 1.19 | 0.583 | 0.396 | 0.161 | 0.559 | 0.174 |
| CD40 | 0.637 | 0.074 | 0.51 | 0.29 | 0.766 | 0.021 | 0.566 | 0.508 |
| CD160 | 0.53 | 0.077 | 0.426 | 0.241 | 0.437 | 0.204 | 0.354 | 0.172 |
| Chemerin | 5.75 | 0.686 | 3.399 | 1.505 | 3.947 | 1.48 | 2.737 | 1.438 |
| Chitinase 3-like 1 | 0.293 | 0.057 | 0.223 | 0.115 | 0.174 | 0.065 | 0.219 | 0.094 |
| Coagulation Factor III | 0.762 | 0.247 | 0.553 | 0.434 | 0.459 | 0.231 | 0.359 | 0.262 |
| Complement Component C5 | 0.172 | 0.043 | 0.146 | 0.094 | 0.121 | 0.051 | 0.099 | 0.052 |
| Complement Factor D | 0.4 | 0.101 | 0.321 | 0.205 | 0.254 | 0.113 | 0.214 | 0.161 |
| C-reactive protein | 0.032 | 0.028 | 0.023 | 0.042 | 7.99E-05 | 0.006 | 0.005 | 0.014 |
| CX3CL1 | 1.045 | 0.206 | 1.167 | 0.649 | 0.781 | 0.224 | 0.777 | 0.391 |
| CXCL1 | 0.516 | 0.084 | 5.548 | 3.496 | 0.795 | 0.089 | 2.018 | 0.435 |
| CXCL2 | 0.446 | 0.119 | 16.595 | 8.854 | 0.404 | 0.054 | 7.588 | 1.604 |
| CXCL9 | 0.506 | 0.113 | 0.416 | 0.269 | 0.529 | 0.408 | 0.283 | 0.195 |
| CXCL10 | 1.396 | 0.446 | 2.344 | 1.066 | 1.162 | 0.677 | 1.572 | 0.634 |
| CXCL11 | 0.422 | 0.082 | 0.319 | 0.155 | 0.279 | 0.125 | 0.259 | 0.189 |
| CXCL13 | 0.45 | 0.084 | 0.347 | 0.168 | 0.289 | 0.106 | 0.266 | 0.141 |
| CXCL16 | 3.173 | 0.757 | 4.347 | 2.103 | 2.057 | 0.873 | 3.167 | 1.645 |
| Cystatin C | 3.788 | 0.631 | 2.968 | 1.466 | 2.576 | 1.156 | 2.298 | 1.422 |
| DKK1 | 0.67 | 0.168 | 0.521 | 0.33 | 0.373 | 0.167 | 0.372 | 0.239 |
| DPPIV | 0.615 | 0.185 | 0.52 | 0.421 | 0.339 | 0.177 | 0.316 | 0.249 |
| EGF | 0.451 | 0.094 | 0.36 | 0.236 | 0.269 | 0.143 | 0.235 | 0.154 |
| Endoglin | 0.129 | 0.068 | 0.085 | 0.077 | 0.074 | 0.033 | 0.068 | 0.061 |
| Endostatin | 2.022 | 0.566 | 1.586 | 0.783 | 1.388 | 0.45 | 1.214 | 0.746 |
| Fetuin A | 0.617 | 0.13 | 0.444 | 0.234 | 0.626 | 0.286 | 0.391 | 0.262 |
| FGF acidic | 0.235 | 0.037 | 0.229 | 0.082 | 0.203 | 0.088 | 0.201 | 0.119 |
| FGF-21 | 0.671 | 0.108 | 0.486 | 0.268 | 0.482 | 0.238 | 0.379 | 0.26 |
| Flt-3 ligand | 0.455 | 0.092 | 0.346 | 0.21 | 0.338 | 0.129 | 0.238 | 0.189 |
| Gas 6 | 0.711 | 0.255 | 0.441 | 0.247 | 0.368 | 0.174 | 0.337 | 0.277 |
| G-CSF | 0.159 | 0.065 | 1.16 | 0.528 | 0.079 | 0.021 | 0.318 | 0.206 |
| GDF-15 | 1.16 | 0.426 | 0.803 | 0.475 | 0.598 | 0.278 | 0.651 | 0.472 |
| GM-CSF | 0.181 | 0.071 | 0.229 | 0.131 | 0.085 | 0.039 | 0.121 | 0.048 |
| HGF | 0.518 | 0.174 | 0.408 | 0.275 | 0.249 | 0.128 | 0.283 | 0.19 |
| ICAM-1 | 0.341 | 0.079 | 0.294 | 0.168 | 0.241 | 0.102 | 0.189 | 0.105 |
| IFN-g | 0.458 | 0.175 | 0.346 | 0.24 | 0.323 | 0.156 | 0.259 | 0.191 |
| IGFBP-1 | 0.252 | 0.065 | 0.163 | 0.093 | 0.161 | 0.081 | 0.131 | 0.092 |
| IGFBP-2 | 0.294 | 0.063 | 0.215 | 0.117 | 0.19 | 0.059 | 0.186 | 0.14 |
| IGFBP-3 | 0.315 | 0.01 | 0.251 | 0.109 | 0.253 | 0.077 | 0.194 | 0.084 |
| IGFBP-5 | 0.566 | 0.132 | 0.429 | 0.246 | 0.402 | 0.18 | 0.296 | 0.17 |

| | LCV uninfected Supernatants | | LCV Infected Supernatants, 24 hr | | SCV Uninfected Supernatants | | SCV Infected Supernatants, 24 hr | |
|---|---|---|---|---|---|---|---|---|
| Secreted Protein | Mean Signal | Standard Deviation | Mean Signal | Standard Deviation | Mean Signal | Standard Deviation | Mean Signal | Standard Deviation |
| IGFBP-6 | 15.539 | 2.677 | 10.383 | 3.831 | 10.504 | 4.082 | 9.059 | 5.396 |
| IL-1a | 0.517 | 0.128 | 0.367 | 0.2 | 0.329 | 0.177 | 0.285 | 0.191 |
| IL-1b | 0.165 | 0.06 | 0.127 | 0.086 | 0.084 | 0.05 | 0.101 | 0.082 |
| IL-1ra | 0.571 | 0.168 | 1.542 | 0.862 | 0.373 | 0.142 | 0.55 | 0.197 |
| IL-2 | 0.06 | 0.042 | 0.03 | 0.04 | 0.005 | 0.014 | 0.014 | 0.021 |
| Il-3 | 0.055 | 0.038 | 0.032 | 0.047 | 0.008 | 0.015 | 0.014 | 0.02 |
| IL-4 | 0.487 | 0.129 | 0.394 | 0.256 | 0.305 | 0.136 | 0.254 | 0.158 |
| IL-5 | 0.345 | 0.138 | 0.218 | 0.157 | 0.19 | 0.089 | 0.153 | 0.108 |
| IL-6 | 0.06 | 0.044 | 2.205 | 1.653 | 0.021 | 0.009 | 0.387 | 0.361 |
| IL-7 | 0.665 | 0.282 | 0.657 | 0.654 | 0.757 | 0.349 | 0.712 | 0.846 |
| IL-10 | 0.688 | 0.117 | 0.551 | 0.347 | 0.482 | 0.229 | 0.376 | 0.269 |
| IL-11 | 0.628 | 0.15 | 0.502 | 0.302 | 0.454 | 0.225 | 0.386 | 0.303 |
| IL-12 p40 | 0.465 | 0.163 | 3.026 | 1.557 | 0.369 | 0.118 | 1.136 | 0.432 |
| IL-13 | 0.356 | 0.135 | 0.361 | 0.334 | 0.217 | 0.119 | 0.228 | 0.205 |
| IL-15 | 0.72 | 0.203 | 0.767 | 0.749 | 0.528 | 0.398 | 0.474 | 0.438 |
| IL-17A | 0.056 | 0.043 | 0.033 | 0.03 | 0.018 | 0.015 | 0.031 | 0.032 |
| IL-22 | 0.33 | 0.096 | 0.234 | 0.131 | 0.17 | 0.087 | 0.172 | 0.11 |
| IL-23 | 0.462 | 0.129 | 0.402 | 0.289 | 0.258 | 0.126 | 0.227 | 0.117 |
| IL-27 p28 | 0.423 | 0.08 | 0.369 | 0.177 | 0.326 | 0.144 | 0.244 | 0.125 |
| IL-28A | 0.794 | 0.248 | 0.577 | 0.322 | 0.586 | 0.256 | 0.438 | 0.332 |
| IL-33 | 0.53 | 0.23 | 0.523 | 0.493 | 0.695 | 0.307 | 0.683 | 0.918 |
| LDL R | 1.499 | 0.954 | 2.867 | 3.975 | 1.887 | 1.172 | 1.052 | 0.92 |
| Leptin | 0.763 | 0.071 | 0.557 | 0.352 | 0.523 | 0.314 | 0.366 | 0.224 |
| LIF | 0.924 | 0.017 | 0.621 | 0.347 | 0.929 | 0.423 | 0.634 | 0.342 |
| Lipocalin-2 | 0.335 | 0.066 | 0.8 | 0.39 | 0.318 | 0.052 | 0.43 | 0.051 |
| LIX | 0.864 | 0.32 | 0.533 | 0.37 | 0.588 | 0.474 | 0.535 | 0.601 |
| M-CSF | 4.13 | 0.941 | 3.126 | 2.096 | 2.731 | 1.188 | 2.205 | 1.857 |
| MMP-2 | 0.603 | 0.142 | 0.453 | 0.257 | 0.403 | 0.208 | 0.314 | 0.207 |
| MMP-3 | 0.14 | 0.074 | 0.132 | 0.1 | 0.06 | 0.031 | 0.074 | 0.047 |
| MMP-9 | 0.103 | 0.053 | 0.535 | 0.271 | 0.036 | 0.02 | 0.181 | 0.057 |
| Myeloperoxidase | 0.835 | 0.272 | 0.633 | 0.475 | 0.437 | 0.2 | 0.567 | 0.313 |
| Osteopontin | 4.974 | 0.7 | 3.707 | 1.991 | 3.349 | 1.535 | 2.849 | 2.065 |
| Osteoprotegrin | 0.159 | 0.078 | 0.194 | 0.219 | 0.125 | 0.05 | 0.103 | 0.093 |
| PD-ECGF | 0.36 | 0.141 | 0.574 | 0.305 | 0.326 | 0.149 | 0.451 | 0.179 |
| PDGF-BB | 0.584 | 0.207 | 0.433 | 0.26 | 0.328 | 0.17 | 0.306 | 0.198 |
| Pentraxin-2 | 0.649 | 0.075 | 0.428 | 0.238 | 0.418 | 0.143 | 0.339 | 0.256 |
| Pentraxin-3 | 0.873 | 0.267 | 0.448 | 0.291 | 0.56 | 0.271 | 0.352 | 0.299 |
| Periostin | 0.198 | 0.037 | 0.084 | 0.05 | 0.094 | 0.064 | 0.084 | 0.079 |
| Pref-1 | 1.314 | 0.34 | 0.811 | 0.472 | 0.765 | 0.398 | 0.609 | 0.436 |
| Proliferin | 0.815 | 0.206 | 0.446 | 0.27 | 0.415 | 0.243 | 0.333 | 0.248 |
| Proprotein Convertase 9 | 0.767 | 0.246 | 0.474 | 0.315 | 0.434 | 0.259 | 0.355 | 0.325 |
| RAGE | 0.74 | 0.307 | 0.423 | 0.265 | 0.321 | 0.161 | 0.326 | 0.235 |
| RBP4 | 0.518 | 0.206 | 0.362 | 0.3 | 0.22 | 0.094 | 0.254 | 0.165 |
| Reg3G | 0.433 | 0.11 | 0.448 | 0.402 | 0.24 | 0.102 | 0.267 | 0.15 |
| Resistin | 0.178 | 0.077 | 0.226 | 0.211 | 0.147 | 0.021 | 0.127 | 0.078 |
| E-Selectin | 0.195 | 0.024 | 0.122 | 0.039 | 0.096 | 0.008 | 0.143 | 0.066 |
| P-Selectin | 0.555 | 0.009 | 0.411 | 0.191 | 0.387 | 0.141 | 0.282 | 0.158 |
| Serpin E1 | 7.454 | 1.446 | 5.816 | 3.057 | 4.833 | 1.938 | 3.938 | 2.516 |
| Serpin F1 | 1.531 | 0.519 | 0.663 | 0.378 | 0.777 | 0.404 | 0.651 | 0.516 |
| Thrombopoetin | 0.813 | 0.147 | 0.458 | 0.273 | 0.392 | 0.221 | 0.324 | 0.229 |
| TIM-1 | 0.843 | 0.246 | 0.425 | 0.253 | 0.414 | 0.235 | 0.324 | 0.271 |
| TNF-a | 0.837 | 0.355 | 1.526 | 0.866 | 0.341 | 0.162 | 0.605 | 0.184 |
| VCAM-1 | 0.443 | 0.258 | 0.197 | 0.136 | 0.128 | 0.033 | 0.114 | 0.047 |
| VEGF | 5.485 | 1.894 | 3.927 | 2.401 | 2.866 | 1.164 | 2.546 | 1.326 |
| WISP-1 | 0.85 | 0.392 | 0.752 | 0.527 | 0.446 | 0.153 | 0.549 | 0.309 |
